# Profound functional and molecular diversity of mitochondria revealed by cell type-specific profiling in vivo

**DOI:** 10.1101/403774

**Authors:** Caroline Fecher, Laura Trovò, Stephan A. Müller, Nicolas Snaidero, Jennifer Wettmarshausen, Sylvia Heink, Oskar Ortiz, Ingrid Wagner, Ralf Kühn, Jana Hartmann, Rosa Maria Karl, Arthur Konnerth, Thomas Korn, Wolfgang Wurst, Doron Merkler, Stefan F. Lichtenthaler, Fabiana Perocchi, Thomas Misgeld

**Author notes:** Current address: Merck KGaA, Darmstadt, Germany. Current address: Max-Delbrück-Centrum für Molekulare Medizin, Berlin, Germany and Berlin Institute of Health, Berlin, Germany. Equal contribution: Caroline Fecher, Laura Trovò. Correspondence to: Thomas Misgeld.

## Abstract

Mitochondria vary in morphology and function in different tissues, however little is known about their molecular diversity among cell types. To investigate mitochondrial diversity in vivo, we developed an efficient protocol to isolate cell type-specific mitochondria based on a new *MitoTag* mouse. We profiled the mitochondrial proteome of three major neural cell types in cerebellum and identified a substantial number of differential mitochondrial markers for these cell types in mice and humans. Based on predictions from these proteomes, we demonstrate that astrocytic mitochondria metabolize long-chain fatty acids more efficiently than neurons. Moreover, we identified Rmdn3 as a major determinant of ER-mitochondria proximity in Purkinje cells. Our novel approach enables exploring mitochondrial diversity on the functional and molecular level in many in vivo contexts.

---

Mitochondria perform different tasks in diverse cell types. For instance, mitochondria control differentiation of immune and stem cells (*1, 2*). Similarly, in the nervous system, mitochondria regulate neurite branching (*3, 4*) and regeneration (*5-7*), as well as synaptic strength, stability and signaling (*8, 9*). While some of these cell type-specific functions are achieved by modulating the well-characterized metabolic roles of mitochondria, also entirely new functions emerge, e.g. in immune or redox signaling (*10-12*), and likely many more remain elusive.

Bulk mitochondrial proteomes vary between tissues (*13*). Nevertheless, it remains unclear to what degree such diversity reflects tissue-specific global differences or variations in cellular composition. Moreover, genes encoding mitochondrial proteins are differentially expressed among tissues, cell types and even cell compartments (*14-16*) – but to what degree such transcriptional diversity results in mitochondrial proteome variability is not resolved. Indeed, while the extrapolation of proteomic data from transcriptomics is generally problematic (*17, 18*), the mitochondrial proteome is uniquely complex, especially in cells of extended geometry, such as neurons (*19*). Hence, in these complex cell types it is crucial to directly measure proteomic variability. Moreover, to test any resulting predictions it is important to obtain functional mitochondria from the respective cell of interest.

Here, we describe such a novel approach that allows isolating functional mitochondria from essentially any cell type. This approach is based on a *MitoTag* mouse, which faithfully expresses an outer mitochondrial membrane (OMM)-targeted green fluorescent protein (GFP) in a Cre recombinase (Cre)-dependent manner. This innocuous and efficient epitope allows capturing cell type-specific mitochondria. We highlight the power of our approach by comparing neural mitochondria of the three major cell populations from cerebellum. Thus, we (i) provide a first systematic profiling of the cell type-specific variability between mitochondrial proteomes in situ; (ii) identify numerous cell type-specific mitochondrial markers in mouse and human brain; (iii) predict and prove metabolic differences in beta-oxidation between neural cells; and (iv) characterize Rmdn3, a cell type-specific mediator of mitochondrial organelle contact sites in cerebellum.

## RESULTS

### *MitoTag* mice allow innocuously tagging of cell type-specific mitochondria in vivo

Previous work established tagging of macromolecular structures like ribosomes to explore cell type-specific mRNA translation profiles in situ (*20, 21*). This approach has been recently extended to organelles, e.g. to explore mitochondrial (*22*) and lysosomal (*23*) metabolomes in vitro, organellar DNA in invertebrates (*24*) or to correlate transcriptional and epigenetic signatures (*25*). To study mitochondrial diversity among cell types in situ, i.e. directly derived from their tissue context, we generated a novel reporter line, named *MitoTag* mouse (**Fig. 1A**). *MitoTag* mice harbor in their Rosa26 locus a floxed “stop” expression cassette, which drives expression of GFP targeted to the outer mitochondrial membrane (OMM) in a Cre-dependent manner (GFP-OMM) (*26*). To achieve GFP-OMM expression in diverse cell types of the central nervous system, *MitoTag* mice were crossed to different Cre-driver lines (**Fig. S1**). Initial characterization was performed on (i) Emx1:Cre crossed to *MitoTag* mice (abbreviated as ‘Emx1:Cre/GFP-OMM’), where the great majority of forebrain neurons and glia contained GFP-OMM-tagged mitochondria (**Fig. 1B**, **S1A**) (*27*); and (ii) on ChAT:Cre/GFP-OMM mice, where mitochondria in cholinergic neurons, including motor neurons, are tagged (**Fig. S1B**). First, we confirmed mitochondrial localization of GFP-OMM by counterstaining Emx1:Cre/GFP-OMM cortex with the endogenous inner mitochondrial membrane protein Cox4 isoform 1 (Cox4i1; **Fig. 1C**). We corroborated OMM localization by expansion microscopy (*28*) in Emx1:Cre/GFP-OMMxThy1:mito-RFP mice (**Fig. 1D**), which additionally express matrix-localized red fluorescent protein (mito-RFP) in neurons. Second, we used ChAT:Cre/GFP-OMM mice to rule out adverse transgene effects on mitochondrial and cellular health in motor neurons by measuring mitochondrial transport and shape, as well as neuromuscular synapse morphology (**Fig. S2A-D**). Finally, crude mitochondrial fractions (CMF) from Emx1:Cre/GFP-OMM cortex showed unchanged oxygen consumption rates compared to controls (**Fig. S2E**).

**Figure 1.**
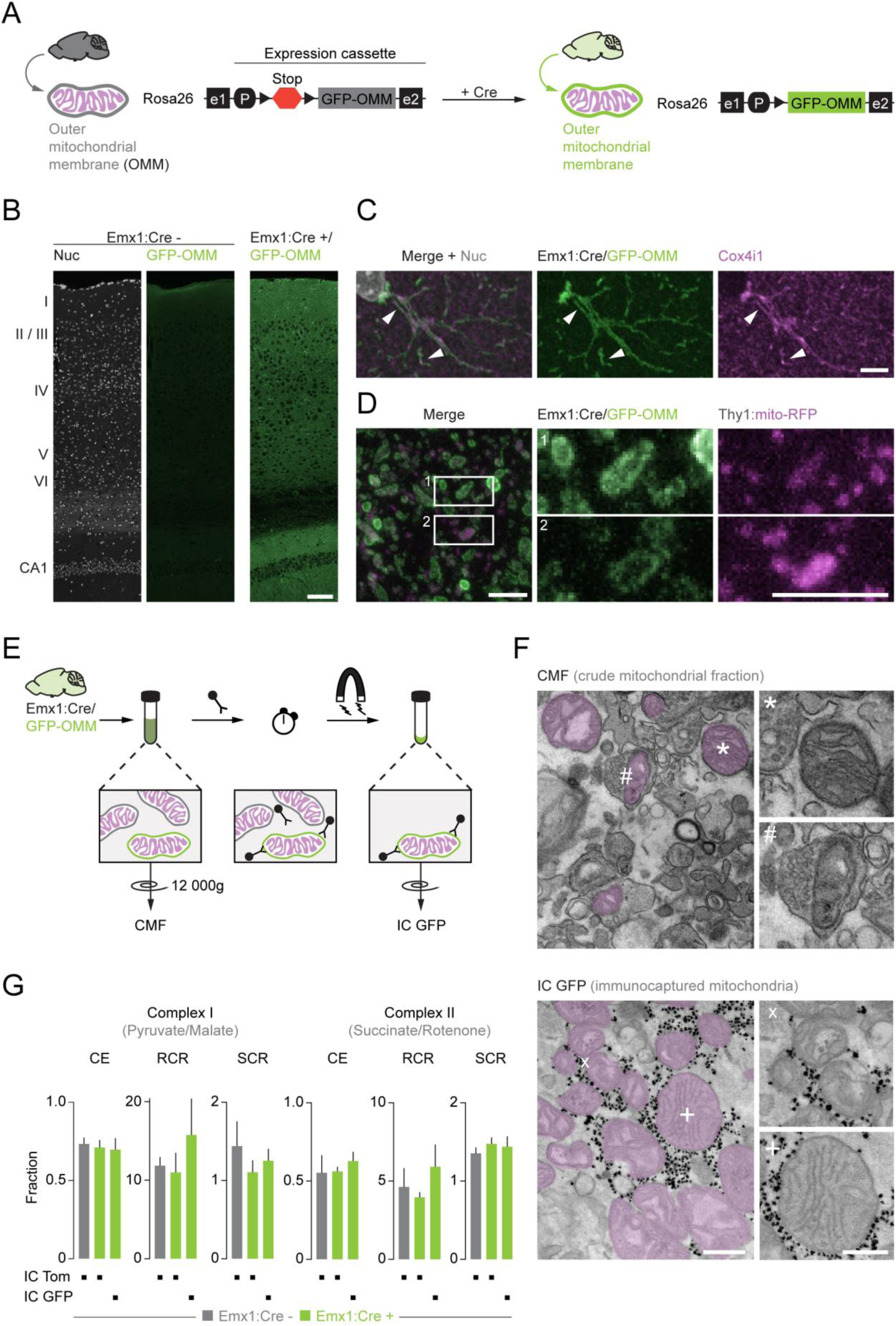
*MitoTag* mice allow isolation of intact and functional mitochondria from cells in situ: (**A**) Schematic of GFP-OMM insertion into the mouse Rosa26 locus (e1/2, exon 1&2; P, promoter). (**B**) Confocal micrographs of cerebral cortex shows no expression in stopped *MitoTag* tissue (left and middle panels; Cre -), but GFP-OMM (green) in neural cells from Emx1:Cre/GFP-OMM mice (right panel; Cre +). Nuclear counterstain, Nuc (left panel; Hoechst, white) shown for layer orientation. (**C**) Confocal imaging of Emx1:Cre/GFP-OMM cerebral cortex immunostained for cytochrome c oxidase subunit 4 isoform 1 (pan-mitochondrial marker, Cox4i1; magenta) confirms mitochondrial localization of GFP-OMM (arrowhead). (**D**) Expansion microscopy of Emx1:Cre/GFP-OMMxThy1:mito-RFP cerebral cortex shows localization of GFP-OMM in the outer mitochondrial membrane (green) surrounding mito-RFP (magenta), which is expressed in the mitochondrial matrix of a large subset of neurons. (**E**) Schematic of the immunocapture (IC) protocol of GFP-OMM tagged mitochondria (CMF, crude mitochondrial fraction). (**F**) Electron micrographs comparing the CMF from Emx1:Cre/GFP-OMM cortex to IC GFP. Note free mitochondria (*) and synaptosome-enclosed mitochondria (#) found in CMF (upper panel) and free, intact mitochondria (x, +) with microbeads (black dots in micrograph, lower panel). (**G**) Bioenergetic parameters derived from oxygen consumption measurements in the presence of pyruvate/malate (left) and succinate/rotenone (right) as substrates for immunocaptured mitochondria. Mitochondria were immunocaptured from Emx1:Cre -/GFP-OMM or Emx1:Cre +/GFP-OMM cortex using Tom22 (pan-mitochondrial, IC Tom) or GFP-OMM (cell type-specific mitochondria, IC GFP) as bait (CE, coupling efficiency; RCR, respiratory capacity ratio; SCR, spare capacity ratio; N=4-7; no significant differences observed with one-way ANOVA with multiple comparisons). Scale bars: 100 µm in B; 5 µm in C, D and inset (note that for D, an estimated expansion factor of ∼2-3 applies); 500 nm in F (250 nm in inset).

To isolate GFP-OMM-tagged mitochondria, we optimized an immunocapture protocol using anti-GFP magnetic microbeads (IC GFP; **Fig. 1E**). This method is well established for mitochondria based on anti-Tom22 microbeads (*29, 30*), but has recently been expanded to neoantigens (*22, 24*). Indeed, for Emx1:Cre/GFP-OMM cortex, Western blot analysis corroborated efficient mitochondrial enrichment via IC GFP (**Fig. S3A**). Using electron microscopy, the majority of GFP-captured objects were isolated mitochondria with well-preserved ultrastructure in contrast to the CMF, which contained synaptosome-enclosed mitochondria and myelin debris (**Fig. 1F**). Moreover, mitochondria that were immunocaptured from either control cortex (Cre^-^ *MitoTag* mice; using anti-Tom22 microbeads, IC Tom) or Emx1:Cre/GFP-OMM cortex (∼75% GFP-OMM-tagged mitochondria; **Fig. S4B**), did not differ in respiratory capacity (**Fig. 1G**).

### *MitoTag* mice allow isolation of cell type-specific mitochondria from complex tissues in situ

A major concern in enriching cell type-specific mitochondria from tissue is that during the isolation, mitochondria from different cells could merge by clumping or fusion (*31*). Indeed, we observed that once we pelleted mitochondria by centrifugation, separation via IC GFP was impaired. Therefore, to determine the selectivity of our immunocapture, we devised “spike-in” experiments with mitochondria carrying mito-RFP. First, we used an ex vivo mixing experiment where cortex from Emx1:Cre/GFP-OMM mice and from Thy1:mito-RFP mice were mixed one-to-one (50:50%) and subjected to immunocapture (**Fig. 2Ai**). Using Western blot analysis, we observed that IC Tom recovered the initial GFP-OMM/mito-RFP ratio of the CMF, while tagged mitochondria were selectively enriched with IC GFP (GFP-OMM: 4.7-fold compared to IC Tom; **Fig. 2B** and **C**). We mimicked cell populations with various abundance by mixing the post-nuclear fractions from Emx1:Cre/GFP-OMM and Thy1:mito-RFP cortex in different ratios and found that the isolation efficacy stayed stable down to dilutions of 1:20 (**Fig. S5A** and **B**). Throughout, we detected ∼ 5% of mito-RFP in IC GFP. To corroborate this experiment in vivo, we used ChAT:Cre/GFP-OMM cortex, where less than 1% of mitochondria are tagged (**Fig. S4B**) – representing a low abundant population of exclusively axonal mitochondria from cholinergic neurons. Still, the immunocapture allowed us to enrich substantial amounts of mitochondria (**Fig. S5C**). Finally, to corroborate enrichment of tagged mitochondria from tissue, we immunocaptured astrocytic mitochondria from Gfap:Cre/GFP-OMMxThy1:mito-RFP cortex (**Fig. 2Aii**). This tissue contained ∼20% GFP-OMM-labeled mitochondria from astrocytes and 63% mito-RFP-positive neuronal mitochondria (**Fig. S4B**). Here, from immediately adjacent cells, GFP-OMM could be enriched to 7.4-fold compared to IC Tom (**Fig. 2D**).

**Figure 2.**
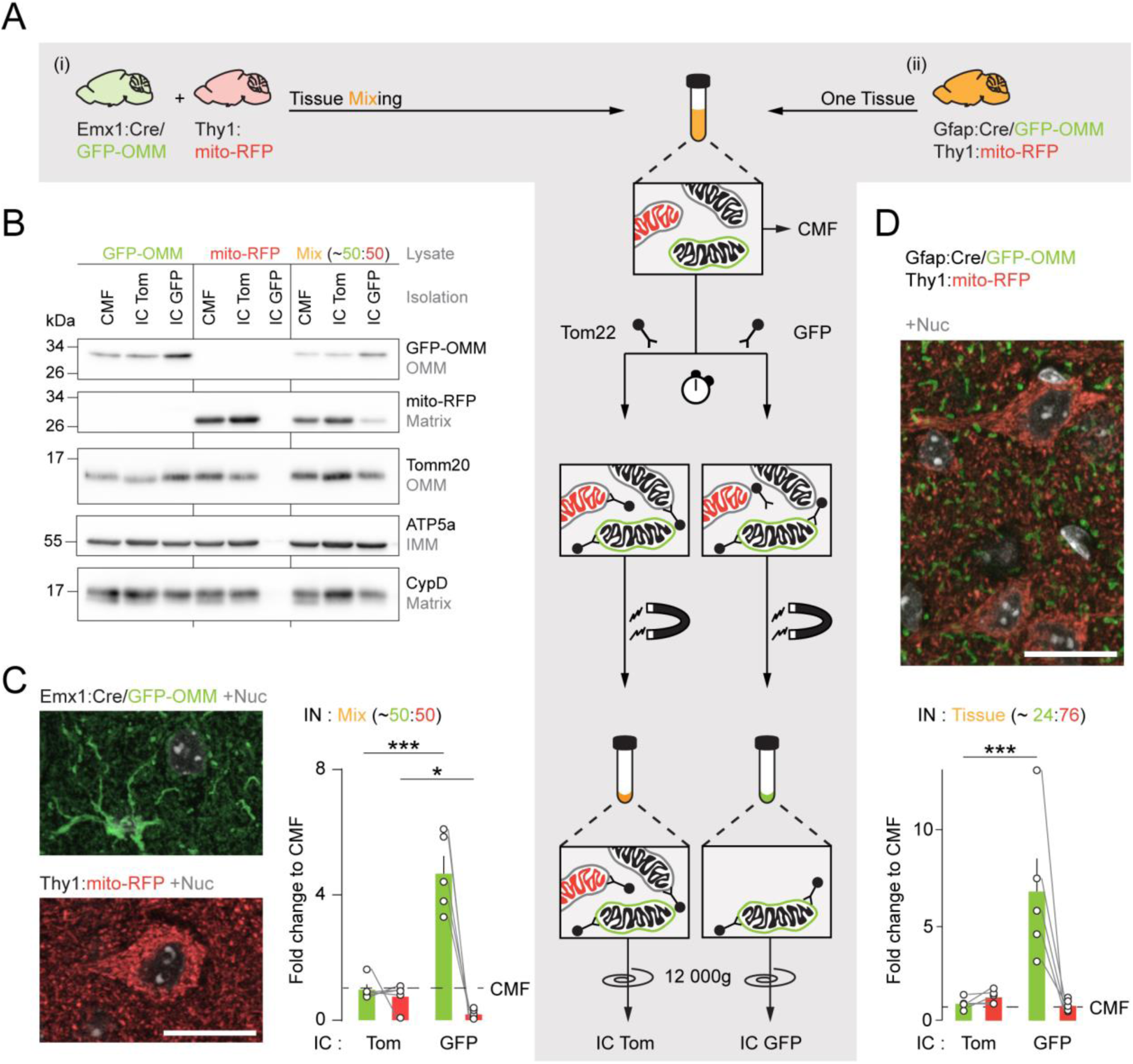
Immunocapture from *MitoTag* mice enriches cell type-specific mitochondria without mitochondrial cross-contamination from other cells: (**A**) Schematic of “spike-in” experiments to demonstrate enrichment of GFP-OMM tagged mitochondria from other mitochondria (mito-RFP). Using cortex, two mitochondrial populations (GFP-OMM, mito-RFP) were mixed either ex vivo on the tissue level (**i**) or in vivo (**ii**) by breeding Gfap:Cre/GFP-OMMxThy1:mito-RFP mice. (**B**, **C**) Enrichment of GFP-OMM-labeled mitochondria after immunocapture via GFP-OMM (IC GFP), but not via Tom22 (IC Tom) from ex vivo mixing experiment (schematic in Ai). Cortex from Emx1:Cre/GFP-OMM (C, top micrograph for tissue expression; GFP-OMM, green; Nuc, white) and cortex from Thy1:mito-RFP (C, bottom micrograph; mito-RFP, red; Nuc, white) were mixed in a 50:50% ratio before lysate generation (B; Mix, orange). (**B**) Western blot analysis of 10 µg protein per condition probed for GFP-OMM, mito-RFP and mitochondrial proteins (OMM, outer mitochondrial membrane – Tomm20; IMM, inner mitochondrial membrane – ATP5a; matrix CypD). Note enrichment of GFP-OMM via IC GFP from pure Emx1:Cre/GFP-OMM tissue (Lysate: GFP-OMM) and the absence of any mitochondrial isolation from pure mito-RFP tissue (Lysate: mito-RFP). (**C**) Quantification of ex vivo mixing experiment (schematic in Ai) comparing isolations to the crude mitochondrial fraction (CMF). Densitometry (GFP-OMM, green bar: p=0.0002; mito-RFP, red bar: p=0.037; N=5, one-tailed ratio-paired *t*-test) is normalized to mitochondrial content. (**D**) In vivo “spike-in” experiment (schematic in Aii) using cortex from Gfap:Cre/GFP-OMMxThy1:mito-RFP mice (micrograph for tissue expression; GFP-OMM, green; mito-RFP, red; Nuc, white) as starting material with a ∼24:76% ratio (cf. Figure S4B). Bar graph shows quantification from Western blot analysis comparing IC Tom and IC GFP to initial CMF (GFP-OMM, p=0.0002; N=5, one-tailed ratio-paired *t*-test). Scale bars: 20 µm in C and D.

### Mitochondrial diversity profiled by label-free proteomics in the adult cerebellum

The cerebellum is composed of various well-characterized cell types, for which specific Cre-driver lines are available, and is prone to mitochondrial dysfunction (*32, 33*). Thus, we used the cerebellum as a proof-of-concept to test whether our *MitoTag* approach could resolve cellular diversity of the mitochondrial proteome in situ. First, we generated *MitoTag* lines for the following cell types (**Fig. 3A**, **S1C**-**E**): (i) Purkinje cells via L7:Cre, the major inhibitory neuron of the cerebellum; (ii) granule cells via Gabra6:Cre, the most abundant excitatory neuron; and (iii) astrocytes via Gfap:Cre. Next, we developed a label-free proteomic workflow for the comparison of mitochondrial proteomes that involved immunocapture of cell type-specific mitochondria (IC GFP) and total tissue mitochondria (IC Tom) per biological replicate (N≥5 mice; **Fig. S6** and **Data Table S1**). IC Tom provided an ‘average mitochondrial proteome’ from cerebellum – thereby correcting for variation among experiments and mouse cohorts. In this background proteome, we quantified 3,718 proteins, which we compared to reference proteomic data sets from mouse cerebellar mitochondria (*13*) and from whole cerebellum (*34*) revealing a coverage of 68% *MitoCarta* (*35*) annotated proteins (**Fig. S3B**). Moreover, intensity-based absolute quantification (iBAQ) demonstrated strong enrichment of mitochondrial proteins over non-mitochondrial proteins (e.g. 6.2-fold for IC TOM from L7:Cre/GFP-OMM) – despite our deliberate choice of a ‘conservative’ isolation approach with the aim to include transiently OMM-associated proteins for discovery and preserve organelle functionality (*36*). In comparison, a recent whole brain proteome study without mitochondrial enrichment revealed that mitochondrial proteins were 1.6-fold more abundant than non-mitochondrial proteins (*34*). In contrast, Percoll-purified mitochondria from cerebellum showed a 6.2-fold increased abundance of mitochondrial over non-mitochondrial proteins in the *MitoCarta* proteomic dataset (*13*) matching our own enrichment. Indeed, 56.5% of peptides accounted for mitochondrial proteins in immunocaptures (**Fig. S3C**). Importantly, mitochondrial proteins showed more prominent abundance changes than non-mitochondrial proteins. For Purkinje cells, 23.1% of mitochondrial annotated proteins showed a significant abundance change (IC GFP/IC Tom ratio >1.5 or <0.67, and p<0.05), while only 4.0% of the non-mitochondrial annotated proteins were significantly changed – suggesting that most of the latter are not specifically captured. We further compared our Purkinje cell-specific dataset (IC GFP of L7:Cre/GFP-OMM; **Fig. S3D**) to the first in vivo whole-cell proteome from Purkinje cells obtained by metabolic labeling of nascent proteins (*37*). This revealed the superior sensitivity of our approach in detecting mitochondrial proteins (*MitoCarta* coverage: 69% vs. 15%).

**Figure 3.**
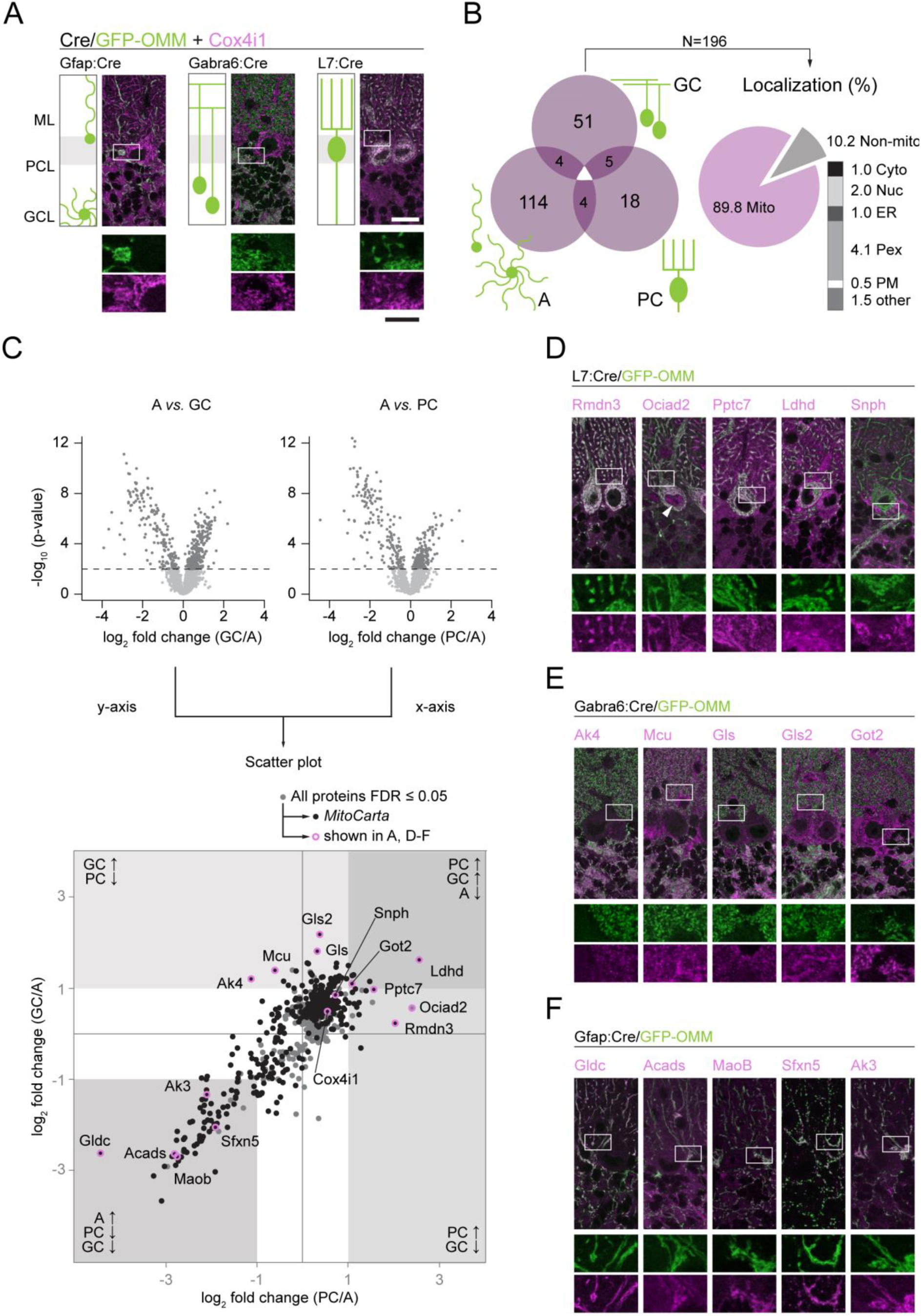
Proteomic profiling of cell type-specific mitochondria in the adult mouse cerebellum: (**A**) Confocal micrographs illustrate GFP-OMM expression patterns (green) and Cox4i1 staining (pan-mitochondrial; magenta) in the investigated cerebellar cell types: astrocytes (left panel, Gfap:Cre; note that molecular layer astrocytes are also known as Bergmann glia), excitatory granule cells (middle panel, Gabra6:Cre) and inhibitory Purkinje cells (right panel, L7:Cre). Each schematic shows cell morphology and anatomical position within the cerebellum (ML, molecular layer; PCL, Purkinje cell layer; GCL, granule cell layer). Details show partial co-localization of cell type-specific GFP-OMM with Cox4i1. (**B**) Venn diagram depicting candidates (N=196) with cell type-specific enrichment obtained by proteomics (fold change ≥│1│ from the pairwise comparison between cell types; PC: Purkinje cells; A: astrocytes; GC: granule cells) and their subcellular localization (Mito, mitochondria; Cyto, cytosol; Nuc, nuclus; ER, endoplasmic reticulum; Pex, peroxisome; PM, plasma membrane; other). (**C**) Comparison of cell type-specific mitochondrial proteomes between cell types (**upper part**: volcano plots comparing GC/A, left; and PC/A, right; two-tailed *t*-test, FDR≤0.05, 500 randomizations, N≥3; dark gray, proteins passing FDR) and among cell types (**lower part**: scatter plot of proteins merging the previous comparison between cell types (FDR≤0.05 in at least one comparison); x-axis, PC/A; y-axis, GC/A). In volcano plots: dark gray circles, proteins FDR≤0.05. In scatter plot: all circles, proteins from volcano plots with FDR≤0.05; black circles, *MitoCarta* annotated proteins; open magenta circles, candidates illustrated in A, D-F. (**D**-**F**) Validation of candidates by immunofluorescence staining in cerebellum. Candidates (magenta) show prominent co-localization with cell type-specific GFP-OMM (green) confirming the proteomic data (see C) in Purkinje cells (**D**), granule cells (**E**) and astrocytes (**F**). Note the pan-neuronal patterns of Ldhd, Gls2 and Got2 (D and E) and the pinceau-like organization of Snph likely in Basket axon collaterals around Purkinje cell soma. GFP-OMM is driven by the Cre-driver lines introduced in A. Arrowhead, nuclear signal. Scale bar: 20 µm in A, and D-F (10 µm in details).

In total, over 85% of identified proteins were shared among all three cell types (**Fig. S6**). We identified 196 candidates with differential expression (log2 fold change ≥│1│ between cell types), of which 89.8% are mitochondrial annotated. Of these, 18, 51 and 114 were exclusively enriched in one cell type (**Fig. 3B**, **S7D**). An additional 13 proteins showed shared enrichment in two cell types. When we compared proteins from astrocytic mitochondria against either mitochondria from granule cell (y-axis; **Fig. 3C**, **S7A** and **D**) or Purkinje cell (x-axis; **Fig. 3C**, **S7B** and **D**), we detected a separation in protein composition of neuronal mitochondria from glial mitochondria. Equally, KEGG pathway (**Fig. S7E**) and GOTerm process analysis (**Fig. S7F**) confirmed differential specializations of neuronal (e.g. ubiquinone biosynthesis) vs. astrocytic mitochondria (e.g. lipid metabolism). In contrast, other mitochondrial protein pathways (e.g. respiratory chain components, protein import and translation, TCA cycle) showed no systematic differences, pointing to a conserved core of the mitochondrial proteome (**Fig. S8**).

We selected 30 candidates for further validation based on literature and antibody availability: Cox4i1 as a ubiquitous mitochondrial marker, syntaphilin (Snph) as a neuronal mitochondrial marker not significantly enriched in the tested cell types and 28 as cell type-enriched proteins. Using immunofluorescence stainings (**Fig. 3D**-**F**, **S9**), we confirmed in all testable cases (23/30) the predicted cell type-specific enrichment and subcellular localization in mitochondria. In seven cases, no confirmation could be achieved due to antibody quality. Some of the validated candidates – especially in astrocytes (**Fig. 3F**) – represent mitochondrial proteins with previously cell type-specific annotation, such as glycine dehydrogenase (Gldc) (*38*), sideroflexin-5 (Sfxn5) (*39*) and monoamine-decarboxylase B (MaoB), which is targeted in positron emission tomography as astrocyte tracer (*40*). Notably, the astrocytic mitochondrial proteome also contained a substantial enrichment of peroxisomal proteins (**Fig. S7E** and **F**, **S9**), some of which are known to have dual targeting (e.g. catalase) (*41, 42*) or tether to mitochondria (e.g. Eci2 and Pex11b) (*43, 44*). In neurons, we expected to identify Snph, a protein involved in regulating mitochondrial distribution and generally believed to be enriched on axonal mitochondria (*45*). Indeed, in line with our proteomics we confirmed only weak signals in Purkinje cell and granule cell mitochondria; however, we additionally observed prominent mitochondrial labeling in Basket axon collaterals synapsing onto Purkinje cell somata – hence forming the pinceau organization (**Fig. 3D**, **S9B**). In some instances, e.g. for mitochondrial adenylate kinase or glutaminase, differential paralogs were detected between cell types (Ak4 vs. Ak3; Gls vs. Gls2; **Fig. 3E** and **F**). In addition, we observed two forms for Ociad2, a non-*MitoCarta* protein dually assigned to mitochondria and endosomes in a recent publication (*46*). A C-terminal antibody confirmed our proteomic prediction of enrichment in Purkinje cell mitochondria, while an N-terminal antibody showed enrichment in granule cells (**Fig. S10**). Surprisingly, we identified cell type-specific enrichment of mitochondrial proteins for which we expected ubiquitous expression – e.g. the mitochondrial calcium uniporter (Mcu; **Fig. 3E**) and its regulators (*47-49*) in granule cells. Similarly, Regulator of microtubule dynamics protein 3 (Rmdn3, also known as PTPIP51), a tether involved in calcium handling (*50*) and lipid transfer (*51*) between the endoplasmic reticulum (ER) and mitochondria, was enriched in Purkinje cells.

Altogether, our validated group of cell type-specific proteins supports the notion that depending on cell type, mitochondria play diverse roles, e.g. in metabolism, calcium handling and organelle communication. Importantly, when we tested a subset of our mitochondrial markers on human cerebellar tissue, we found the expression patterns to be preserved (**Fig. 4**) opening up the possibility to explore cell type-specific mitochondrial pathology in humans. Thus, our results point to a so far unappreciated profound organellar diversity among cell types in vivo.

**Figure 4.**
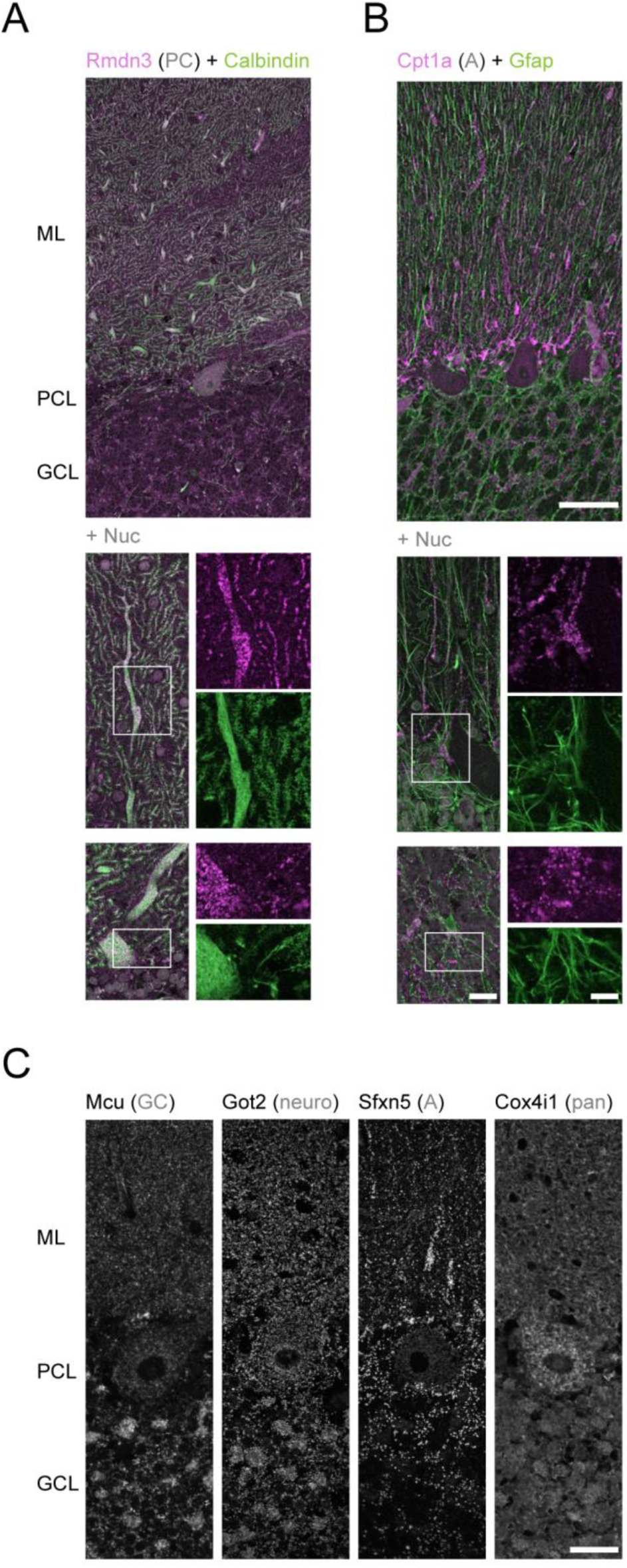
Cell type-specific mitochondrial markers are conserved in human cerebellum: (**A**, **B**) Overview with details of immunostainings for Rmnd3 (**A**, magenta) and Cpt1a (**B**, magenta) in post mortem human cerebellum. (**A**) Rmdn3 marks Purkinje cell mitochondria (PC; cell type marker: calbindin, green). (**B**) Cpt1a identifies astrocytic mitochondria (A; cell type marker Gfap, green). Nuc, nuclear counterstain (DAPI; white). (**C**) Details of four cell type-specific mitochondrial markers illustrate the diverse shapes and quantities of mitochondria in human cerebellum. Mcu is enriched in granule cell (GC; first from left). Note intense signal in granule cell layer in comparison to Purkinje cell soma and molecular layer signals. Got2 is pan-neuronal (neuro; second from left) and Sfxn5 is astrocytic (A; second from right). Cox4i1 shows pan-mitochondrial labeling (pan, first from right). Note the broad signal with loss of individual mitochondrial shapes due to proximity. ML, molecular layer; PCL, Purkinje cell layer; GCL, granule cell layer. Scale bars: 50 µm in A and B upper panel; 20 µm in A and B lower panel (10 µm in details); 25 µm in C.

### Fatty acids are more efficiently metabolized by astrocytic than neuronal mitochondria

Pathway analysis between neuronal and astrocytic mitochondria revealed an enrichment of enzymes involved in mitochondrial beta-oxidation (**Fig. 5A**, **S7E**). However, the brain is often assumed to rely on glucose and lactate for energy production, rather than lipids (*52, 53*). Still, neural cells appear to respire on fatty acids under certain conditions (*54, 55*) with important functional implications, e.g. stem cell proliferation (*56, 57*). Despite this, whether long-chain fatty acids support respiration in neural cells under steady-state conditions remains controversial. Guided by our proteomic profiling, we addressed this question by first confirming the astrocytic enrichment of two enzymes involved in beta-oxidation by immunofluorescence staining, short-chain specific acyl-CoA dehydrogenase (Acads; **Fig. 3F**, **S9C**) and carnitine palmitoyltransferase 1a (Cpt1a; **Fig. 5B**), a rate limiting enzyme in long-chain fatty acids oxidation (*58*). Secondly, we used our approach to compare oxygen consumption rates of in situ isolated cell type-specific mitochondria on the C16-fatty acid, palmitoyl-carnitine (**Fig. 5C**, **S11**). We chose a substrate that is preferentially metabolized in mitochondria due to its carbon chain length and carnitine pre-conjugation (*59, 60*), because astrocytic mitochondria appear to engage selectively with peroxisomes based on our proteomic profiling (**Fig. S7E** and **F**). Astrocytic mitochondria performed significantly better than Purkinje cell-derived mitochondria in coupling efficiency (astrocytic vs. Purkinje cell mitochondria; CE:1.2-fold; p=0.03, one-tailed Welch’s t-test; N≥9), respiratory capacity ratio (RCR: 1.6-fold; p<0.001) and spare capacity ratio (SCR: 1.7-fold; p<0.001). In contrast, selective respiration via complex I and II did not differ significantly (**Fig. 5D** and **E**, **S11**).

**Figure 5.**
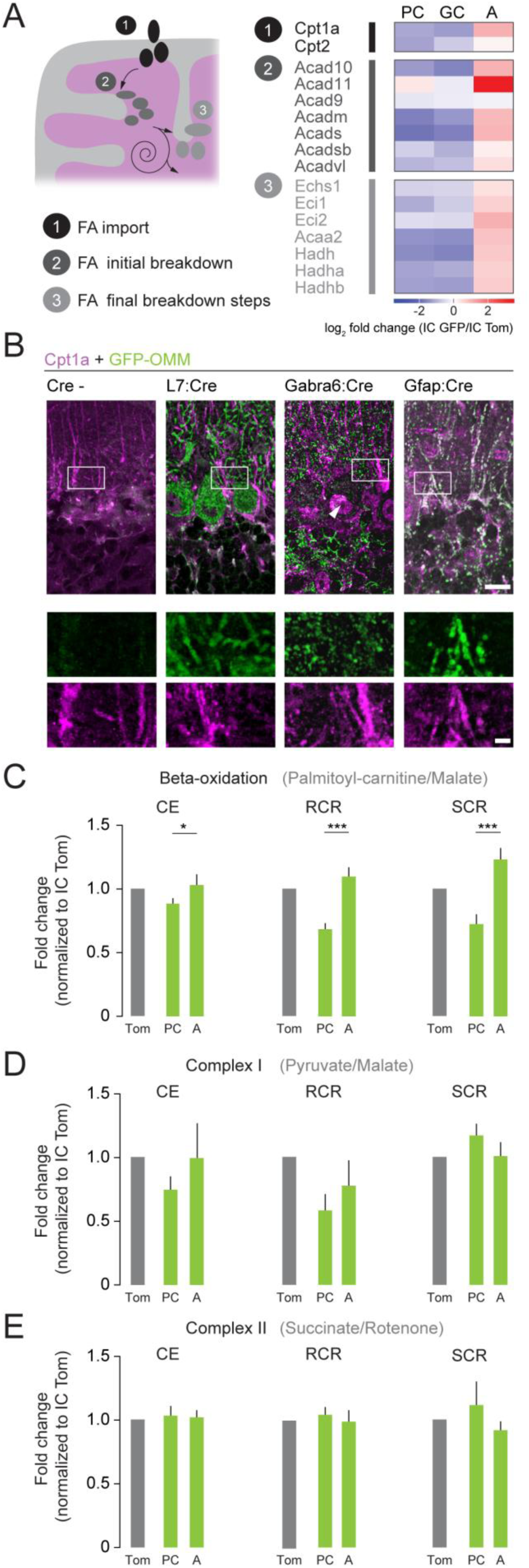
Astrocytic mitochondria metabolize long-chain fatty acids more efficiently than neurons: **(A)** Schematic depicting the localization of the three steps of beta-oxidation in mitochondria: (A) Fatty acid (FA) import, (2) FA initial breakdown and (3) FA final breakdown. Heat map showing cell type-specific protein changes for enzymes involved in beta-oxidation as average log_2_ fold change of IC GFP/IC Tom (PC, Purkinje cell; GC, granule cell; A, astrocyte).(B) Immunostaining for carnitine palmitoyltransferase 1a (Cpt1a, magenta) co-localizes with astrocytic mitochondria in cerebellum (right panel; Gfap:Cre/GFP-OMM, green). Staining is additionally shown in tissue from Cre -, L7:Cre/GFP-OMM and Gabra6:Cre/GFP-OMM mice. Arrowhead, nuclear signal as previously reported for Cpt1a (*80*). (**C**-**E**) Bioenergetic parameters derived from oxygen consumption measurements of immunocaptured mitochondria from Purkinje cells (PC, L7:Cre/GFP-OMM) or astrocytes (A, Gfap:Cre/GFP-OMM), in the presence of different substrates (CE, coupling efficiency; RCR, respiratory capacity ratio; SCR, spare capacity ratio). (**C**) All parameters under palmitoyl-carnitine/malate are significantly higher in astrocytic mitochondria (CE: p=0.0436; RCR: p=0.0001; SRC: p<0.0001, N=9-10; unpaired *t*-test with Welch’s correction). (**D**) Parameters under pyruvate/malate and (**E**) succinate/rotenone do not show any significant difference. For comparison, parameters are normalized to corresponding IC Tom value (gray bar; cf. Figure S11 for non-normalized values). Scale bar: 25 µm in C (5 µm in detail).

### Rmdn3 mediates ER-mitochondria juxtapositions in Purkinje cells

A central aspect of mitochondrial function is communication with other organelles, e.g. via contact sites (*61-63*). Rmdn3 is an established tether between the ER and mitochondria involved in calcium and lipid transfer (*50*). In our proteomic profiling, we found this OMM protein (*64*) to be enriched 3.5-fold over granule cell mitochondria and 4.1-fold over astrocytic mitochondria. Previous reports established the interaction of Rmdn3 with ER-membrane proteins, namely VAPB (an amyotrophic lateral sclerosis-associated protein) (*50*) and ORP5/8 (*51*). First, we confirmed by immunofluorescence staining that Rmdn3 is highly enriched in Purkinje cell mitochondria (**Fig. 6A**) (*65*). Next, we tested the resulting prediction of more extensive mitochondria-ER contacts in Purkinje cells compared to astrocytes by ultrastructural analysis. Indeed, ER-mitochondria contacts were 2.7-fold more extensive in Purkinje cell somata compared to neighboring astrocytes (≤30 nm distance: 38% Purkinje cells vs. 14% astrocytes; two-tailed unpaired *t*-test, p<0.0001, N≥10 cells from two mice; **Fig. 6B**, **S13A**). Finally, we confirm that these juxtapositions are affected by Rmdn3 levels using a new knockout mouse model. These mice showed a gene dose-dependent protein loss in cerebellum by Western blot and immunofluorescence analysis (**Fig. 6C** and **D**), even though some non-specific bands and residual non-neuronal staining persisted. Ultrastructural appraisal demonstrated a reduction of ER-mitochondria contacts in Purkinje cell somata (+/+, 37%; +/-, 28%; -/-, 20%; one-way ANOVA with Tukey’s post-hoc test, +/+ vs. -/-: p<0.0001 and +/+ vs. +/-: p=0.0022, +/- vs. -/-: p=0.0016, N≥8 cells from 2-3 mice; **Fig. 6E**, **S13B**), while contacts in astrocytes were not significantly altered.

**Figure 6.**
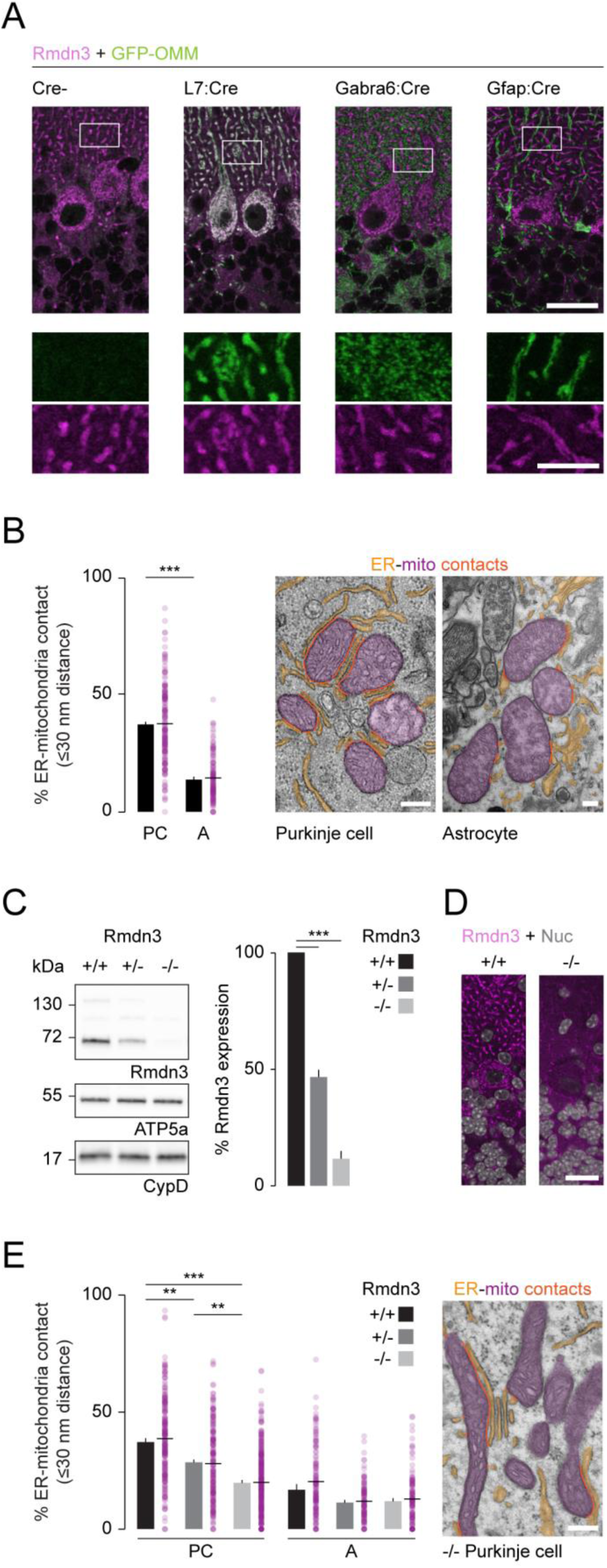
Rmdn3 mediates close ER-mitochondria juxtapositions in Purkinje cells: **(A)** Immunostaining for Rmdn3 (magenta) co-localizes with Purkinje cell mitochondria in mouse cerebellum (second panel from left; L7:Cre/GFP-OMM, green). Staining is additionally shown in tissue from Cre-, Gfap:Cre/GFP-OMM and Gabra6:Cre/GFP-OMM mice. (B) Ultrastructural analysis of ER-mitochondria proximity in wild type cerebellum reveals the degree of proximity of ER (orange) to mitochondria (magenta) in the cytoplasm of Purkinje cells (left panel) and adjoining astrocytes (right panel). Note differing scale bar sizes due to mitochondrial size. The contact frequency between organelles is shown as % mitochondrial perimeter in distance ≤30 nm (red lines in panels) and illustrated as bar graph using cells as statistical entities (p<0.0001, N= 10 vs. 16 cells from two animals; two-tailed unpaired *t*-test). Dot plot shows the population of mitochondria per genotype and cell type (N≥120 mitochondria, magenta dots, from two mice). (**C**, **D**) Rmdn3 knockout mice show a gene dose-dependent reduction in Rmdn3 protein levels in the cerebellum. (**C**) 20 µg protein from cerebellum of wild type (+/+), heterozygous (+/-) and knockout (-/-) mice were analyzed via Western blot probing for Rmdn3 (size: 51 and >130 kDa) and mitochondrial proteins (ATP5a, CypD). Bar graph shows quantification of Rmdn3 levels relative to mitochondrial content (+/+ vs. -/-: p<0.0001, +/+ vs. +/-: p=0.0003, N=4; two-tailed paired t-test).(**D**) Immunostaining for Rmdn3 (magenta) in wild type (left panel; +/+) and knockout (right panel; -/-) cerebellum shows loss of mitochondrial pattern in Purkinje cells. Weak residual signal in astrocytic mitochondria might be due to imperfect specificity of the antibody for the canonical form (52 kDa) of Rmdn3, cf. Figure S12B. Nuclei, nuc, white. (**E**) Ultrastructural analysis of ER-mitochondria proximity in Rmdn3 knockout cerebellum reveals reduced contact frequency (ER, orange; mitochondria, magenta; contact, red lines) in the cytoplasm of Purkinje cells (electron micrograph). Bar graph shows the decrease in contact frequency between organelles based on cells as statistical entities (≤30 nm distance; +/+ vs. -/-: p<0.0001, +/+ vs.+/-: p=0.0022, +/- vs. -/-: p=0.0016, N≥8 cells from 2-3 mice; one-way ANOVA with Tukey’s post-hoc test) in Purkinje cells but not in astrocytes (+/+ vs. -/-: p=0.17, +/+ vs. +/-: p=0.12, N≥10 cells from 2-3 mice). Dot plot shows the population of mitochondria per genotype and cell type (N≥105 mitochondria, magenta dots, from 2-3 mice). Scale bars: 25 µm in A (5 µm in detail); 200 nm in B, E; 20 µm in D.

## DISCUSSION

Here, we describe a versatile approach to obtain intact cell type-specific mitochondria from complex tissues. This approach allows for functional and ‘omics’-level analysis of mitochondria derived from any cell type during development, physiology and disease. We demonstrate this by comparing the mitochondrial proteomes of the principal inhibitory and excitatory neurons, as well as of the major glial cell type in the mouse cerebellum.

Compared to other methods that explore the molecular composition of mitochondria, our approach has a number of distinct advantages: (i) Mice of all ages and genotypes can be used, enabling studies of aging-related changes or disease models, which for many cell types are not possible with cell culture- or tissue dissociation-based approaches. (ii) In cells with extended morphology, mitochondria from all compartments can be obtained. We illustrate this by capturing axonal mitochondria from cholinergic forebrain projections (**Fig. S4B, S5C**). (iii) While transcriptomes provide insights into the regulation of mitochondrial gene expression, mitochondrial proteomes are shaped by multiple layers of regulation (*19, 66, 67*) and hence are even less predictable from transcriptomes than global cellular proteomes (*17, 18*) – a limitation that our direct proteomic approach circumnavigates. (iv) Our approach can yield proteins that are uncharacterized or only transiently associated with mitochondria that would be otherwise overlooked by methods relying on prior knowledge of mitochondrial localization. (v) In contrast to methods that are based on the affinity-purification of labeled proteins from cell types, e.g. via BONCAT (*37*) or APEX *(68*), our approach provides the intact and functional organelle for correlated analysis of the mitochondrial genome or transcriptome (*14, 24*) as well as of mitochondrial functions, e.g. respiration, calcium handling or ROS production. This is further aided by the fluorescent tag, which allows in situ observation, and the bulk IP approach, which is relatively swift and gentle – an important feature for functional and metabolic investigations (*22*).

Indeed, when applied to the adult cerebellum, our approach yielded a number of novel insights: (i) It revealed profound mitochondrial diversity among cell types, with a significant part of the annotated mitochondrial proteome (∼ 15%) being differentially regulated among the cell types we investigated. (ii) Our verified cell type-specific mitochondrial proteins provide a set of in situ markers to characterize the number, distribution and morphology of mitochondria in specific cerebellar cell types without genetic labels across developmental stages and disease models. Several of these new mitochondrial markers are conserved in human tissue (**Fig. 4**), thus opening the path towards characterizing cell type-resolved mitochondrial changes in human disease. So far, this has only been possible for cellular compartments that are large enough to be resolved by light microscopy (*69*) or by using electron microscopy, which is cumbersome and highly susceptible to post mortem changes. (iii) Pathway analysis of our cerebellar proteomes resulted in clear predictions for differentially regulated metabolic pathways (**Fig. S7E**), which we could directly confirm by demonstrating efficient beta-oxidation in immunocaptured astrocytic mitochondria (**Fig. 5C**, **S11**). (iv) Chiming with this result, we observed an enrichment of peroxisomal proteins in captured mitochondria from astrocytes (**Fig. S7E** and **F**). As peroxisomes are another important compartment for lipid metabolism (*70*) and engage intimately with mitochondria, this interaction might enhance glial use of lipids and the exchange of proteins between mitochondria and peroxisomes, *e.g.* as part of peroxisome biogenesis (*71*). (v) Our proteomic profiling predicted candidates that mediate cell type-specific regulation of organelle contacts. For instance, we found a recently identified mitochondria-peroxisome tether, Pex11B (*44*), enriched in astrocytic mitochondria. Similarly, we identified Rmdn3 enriched in Purkinje cell mitochondria, which mediates ER-mitochondria contact sides (**Fig. 6**). (vi) The enrichment of Rmdn3 fits well with independent observations indicating unique intracellular calcium handling mechanisms in Purkinje cells relating to the distinct morphology, firing pattern and synaptic plasticity of this neuron (*72*). In contrast, mitochondrial calcium handling in granule cells seems to be governed by strong expression of the Mcu complex (**Fig. 3E**). A similar regulation of mitochondrial calcium handling has been observed in other excitable tissues (*48, 73-75*) either by tissue-specific expression of Mcu components (Micu1-3) or by posttranslational modification (e.g. redox state) (*76*). The low expression of the Mcu complex in Purkinje cells thus matches the close ER-mitochondria proximity mediated by Rmdn3, but could also relate to the high levels of cytoplasmic calcium buffers in Purkinje cells (e.g. parvalbumin), which in other cells downregulate Mcu expression (*77*). Overall this hints at hitherto unappreciated cell type-specific regulation of even basic mitochondrial functions in the central nervous system (*78*).

Our data reveal that underneath the initial perception of mitochondria as rather uniform and autonomous organelles lies a layer of profound cell type-specific mitochondrial biology that we are only starting to appreciate. Given the startling tissue- and cell type-specificity of many pathologies involving mitochondria (*2, 79*) and the increasing number of roles ascribed to mitochondria as signaling hub, the *MitoTag* approach provides a valuable tool to recognize mitochondria fully as the multifaceted organelles that they are.

## Acknowledgments

We thank M. and N. Budak, as well as S. Taskin for animal husbandry; Anna Berghofer, A. Graupner, Y. Hufnagel, K. Wullimann and M. Schetterer for technical and administrative support. We thank U. Fuenfschilling (MPI Goettingen) for providing Gabra6:Cre mice, D. Crane (Griffith University) for PEX14 antibody and N. Mizushima (Addgene plasmid #38249) for the pMXs-IP GFP-Omp25 plasmid. We are grateful to L. Jiang for help with bioinformatics analyses. We thank L. Godinho, as well as M. and O. Schuldiner for reading an earlier version of this manuscript;

## Funding

This work was supported by the Deutsche Forschungsgemeinschaft (DFG) through the Munich Center for Systems Neurology (SyNergy; EXC 1010; support to A.K., T.K., S.L., T.M., F.P., W.W.), the Center for Integrated Protein Science Munich (CIPSM, EXC 114; to A.K. and T.M.), Collaborative Research Centers CRC870 (T.M.), CRC1054 (T.K.) and CRC-TR128 (T.K.), as well as research unit FOR2290 (S.F.L) and research grants Mi694/7-1/8-1 (T.M.). F.P. and J.W. were supported by the Emmy-Noether program of the DFG (Pe2053/1-1 to F.P.). Further support came from the European Research Council (ERC) under the European Union’s Seventh Framework Program (FP/2007-2013; ERC Grant Agreement n. CoG 616791 with T.M. and CoG 647215 to T.K.) and the German Center for Neurodegenerative Diseases (DZNE Munich; to S.L., T.M., W.W.). L.T. was supported by an EMBO Long-Term Fellowship (EMBO ALTF 108-2013). Further support came from the Centers of Excellence in Neurodegeneration and the Helmholtz-Israel Program (to S.F.L), the Swiss National Science Formation and HHS Foundation (to D.M.), the German Federal Ministry of Education and Research (BMBF) through “T-B interaction in NMO” to T.K. and “Mitochondrial endophenotypes of Morbus Parkinson” (MitoPD, 031A430E) to W.W..

## Author contributions

C.F., L.T. and T.M. devised the study. L.T., T.M., O.O., R.K. and W.W. designed and generated the *MitoTag* mice, which C.F. and L.T. characterized. C.F. designed the IC protocol (with support from J.W. and F.P.) and performed most of the isolations. S.M. and S.L. performed sample preparation, mass spectrometry and primary data analysis. C.F. further analyzed the proteomic data sets. L.T. performed bioenergetics measurements and corresponding isolations with support from J.W. and F.P.. C.F., S.H. and T.K. performed and analyzed flow cytometry. C.F. performed Western blot analysis and validation of candidates by immunofluorescence staining. N.S. and C.F. obtained and analyzed electron microscopy data. C.F. characterized the Rmdn3-/- mouse model. J.H., R.M.K. and A.K. provided pilot experiments for single-cell characterization in the cerebellum. I.W. and D.M. performed immunofluorescence staining on human tissue. C.F., L.T., and T.M. wrote the paper with input from all authors.

## Competing interests

The authors declare no competing interests.

## Data and materials availability

Data and material can be requested from the corresponding author upon reasonable request.

## References

1. N. J. MacIver, R. D. Michalek, J. C. Rathmell, Metabolic regulation of t lymphocytes. Annu Rev Immunol 31, 259–283 (2013).

2. H. Chen, D. C. Chan, Mitochondrial dynamics in regulating the unique phenotypes of cancer and stem cells. Cell Metab 26, 39–48 (2017).

3. M. Spillane, A. Ketschek, T. T. Merianda, J. L. Twiss, G. Gallo, Mitochondria coordinate sites of axon branching through localized intra-axonal protein synthesis. Cell Rep 5, 1564–1575 (2013).

4. J. Courchet et al., Terminal axon branching is regulated by the lkb1-nuak1 kinase pathway via presynaptic mitochondrial capture. Cell 153, 1510–1525 (2013).

5. R. Cartoni et al., The mammalian-specific protein armcx1 regulates mitochondrial transport during axon regeneration. Neuron 92, 1294–1307 (2016).

6. S. M. Han, H. S. Baig, M. Hammarlund, Mitochondria localize to injured axons to support regeneration. Neuron 92, 1308–1323 (2016).

7. B. Zhou et al., Facilitation of axon regeneration by enhancing mitochondrial transport and rescuing energy deficits. J Cell Biol 214, 103–119 (2016).

8. M. J. Devine, J. T. Kittler, Mitochondria at the neuronal presynapse in health and disease. Nat Rev Neurosci 19, 63–80 (2018).

9. B. C. Yoon et al., Local translation of extranuclear lamin b promotes axon maintenance. Cell 148, 752–764 (2012).

10. J. Nunnari, A. Suomalainen, Mitochondria: In sickness and in health. Cell 148, 1145–1159 (2012).

11. A. P. West, G. S. Shadel, S. Ghosh, Mitochondria in innate immune responses. Nat Rev Immunol 11, 389–402 (2011).

12. H. M. McBride, M. Neuspiel, S. Wasiak, Mitochondria: More than just a powerhouse. Curr Biol 16, R551–560 (2006).

13. D. J. Pagliarini et al., A mitochondrial protein compendium elucidates complex i disease biology. Cell 134, 112–123 (2008).

14. T. R. Mercer et al., The human mitochondrial transcriptome. Cell 146, 645–658 (2011).

15. T. Shigeoka et al., Dynamic axonal translation in developing and mature visual circuits. Cell 166, 181–192 (2016).

16. Y. Zhang et al., An rna-sequencing transcriptome and splicing database of glia, neurons, and vascular cells of the cerebral cortex. J Neurosci 34, 11929–11947 (2014).

17. A. Ghazalpour et al., Comparative analysis of proteome and transcriptome variation in mouse. PLoS Genet 7, e1001393 (2011).

18. Z. Khan et al., Primate transcript and protein expression levels evolve under compensatory selection pressures. Science 342, 1100–1104 (2013).

19. T. Misgeld, T. L. Schwarz, Mitostasis in neurons: Maintaining mitochondria in an extended cellular architecture. Neuron 96, 651–666 (2017).

20. E. Sanz et al., Cell-type-specific isolation of ribosome-associated mrna from complex tissues. Proc Natl Acad Sci U S A 106, 13939–13944 (2009).

21. M. Heiman et al., A translational profiling approach for the molecular characterization of cns cell types. Cell 135, 738–748 (2008).

22. W. W. Chen, E. Freinkman, T. Wang, K. Birsoy, D. M. Sabatini, Absolute quantification of matrix metabolites reveals the dynamics of mitochondrial metabolism. Cell 166, 1324–1337.e1311 (2016).

23. M. Abu-Remaileh et al., Lysosomal metabolomics reveals v-atpase- and mtor-dependent regulation of amino acid efflux from lysosomes. Science 358, 807–813 (2017).

24. A. Ahier et al., Affinity purification of cell-specific mitochondria from whole animals resolves patterns of genetic mosaicism. Nat Cell Biol 20, pp. 352–360 (2018).

25. H. C. Roh et al., Simultaneous transcriptional and epigenomic profiling from specific cell types within heterogeneous tissues in vivo. Cell Rep 18, 1048–1061 (2017).

26. C. Horie, H. Suzuki, M. Sakaguchi, K. Mihara, Characterization of signal that directs c-tail-anchored proteins to mammalian mitochondrial outer membrane. Mol Biol Cell 13, 1615–1625 (2002).

27. J. A. Gorski et al., Cortical excitatory neurons and glia, but not gabaergic neurons, are produced in the emx1-expressing lineage. J Neurosci 22, 6309–6314 (2002).

28. P. W. Tillberg et al., Protein-retention expansion microscopy of cells and tissues labeled using standard fluorescent proteins and antibodies. Nat Biotechnol 34, 987–992 (2016).

29. A. Franko et al., Efficient isolation of pure and functional mitochondria from mouse tissues using automated tissue disruption and enrichment with anti-tom22 magnetic beads. PloS one 8, e82392 (2013).

30. H. T. Hornig-Do et al., Isolation of functional pure mitochondria by superparamagnetic microbeads. Anal Biochem 389, 1–5 (2009).

31. S. Meeusen, J. M. McCaffery, J. Nunnari, Mitochondrial fusion intermediates revealed in vitro. Science 305, 1747–1752 (2004).

32. H. Chen, J. M. McCaffery, D. C. Chan, Mitochondrial fusion protects against neurodegeneration in the cerebellum. Cell 130, 548–562 (2007).

33. D. Di Bella et al., Mutations in the mitochondrial protease gene afg3l2 cause dominant hereditary ataxia sca28. Nat Genet 42, 313–321 (2010).

34. K. Sharma et al., Cell type- and brain region-resolved mouse brain proteome. Nat Neurosci 18, 1819–1831 (2015).

35. S. E. Calvo, K. R. Clauser, V. K. Mootha, Mitocarta2.0: An updated inventory of mammalian mitochondrial proteins. Nucleic Acids Res 44, D1251–1257 (2016).

36. X. Wang et al., Isolation of brain mitochondria from neonatal mice. J Neurochem 119, 1253– 1261 (2011).

37. B. Alvarez-Castelao et al., Cell-type-specific metabolic labeling of nascent proteomes in vivo. Nat Biotechnol 35, 1196–1201 (2017).

38. K. Sato, S. Yoshida, K. Fujiwara, K. Tada, M. Tohyama, Glycine cleavage system in astrocytes. Brain Res 567, 64–70 (1991).

39. S. Miyake et al., Identification and characterization of a novel mitochondrial tricarboxylate carrier. Biochem Biophys Res Commun 295, 463–468 (2002).

40. J. S. Fowler et al., Monoamine oxidase: Radiotracer development and human studies. Methods 27, 263–277 (2002).

41. V. Y. Petrova, D. Drescher, A. V. Kujumdzieva, M. J. Schmitt, Dual targeting of yeast catalase a to peroxisomes and mitochondria. Biochem J 380, 393–400 (2004).

42. S. E. Calvo et al., Comparative analysis of mitochondrial n-termini from mouse, human, and yeast. Mol Cell Proteomics 16, 512–523 (2017).

43. J. Fan, X. Li, L. Issop, M. Culty, V. Papadopoulos, Acbd2/eci2-mediated peroxisome-mitochondria interactions in leydig cell steroid biosynthesis. Mol Endocrinol 30, 763–782 (2016).

44. N. Shai et al., Systematic mapping of contact sites reveals tethers and a function for the peroxisome-mitochondria contact. Nat Commun 9, 1761 (2018).

45. J. S. Kang et al., Docking of axonal mitochondria by syntaphilin controls their mobility and affects short-term facilitation. Cell 132, 137–148 (2008).

46. S. Sinha, V. A. Bheemsetty, M. S. Inamdar, A double helical motif in ociad2 is essential for its localization, interactions and stat3 activation. Sci Rep 8, 7362 (2018).

47. J. M. Baughman et al., Integrative genomics identifies mcu as an essential component of the mitochondrial calcium uniporter. Nature 476, 341–345 (2011).

48. D. De Stefani, A. Raffaello, E. Teardo, I. Szabò, R. Rizzuto, A forty-kilodalton protein of the inner membrane is the mitochondrial calcium uniporter. Nature 476, 336–340 (2011).

49. K. J. Kamer, V. K. Mootha, The molecular era of the mitochondrial calcium uniporter. Nat Rev Mol Cell Biol 16, 545–553 (2015).

50. K. J. De Vos et al., Vapb interacts with the mitochondrial protein ptpip51 to regulate calcium homeostasis. Hum Mol Genet 21, 1299–1311 (2012).

51. R. Galmes et al., Orp5/orp8 localize to endoplasmic reticulum-mitochondria contacts and are involved in mitochondrial function. EMBO Rep 17, 800–810 (2016).

52. P. Mergenthaler, U. Lindauer, G. A. Dienel, A. Meisel, Sugar for the brain: The role of glucose in physiological and pathological brain function. Trends Neurosci 36, 587–597 (2013).

53. P. Schönfeld, G. Reiser, Why does brain metabolism not favor burning of fatty acids to provide energy? Reflections on disadvantages of the use of free fatty acids as fuel for brain. J Cereb Blood Flow Metab 33, 1493–1499 (2013).

54. J. Edmond, R. A. Robbins, J. D. Bergstrom, R. A. Cole, J. de Vellis, Capacity for substrate utilization in oxidative metabolism by neurons, astrocytes, and oligodendrocytes from developing brain in primary culture. J Neurosci Res 18, 551–561 (1987).

55. D. Ebert, R. G. Haller, M. E. Walton, Energy contribution of octanoate to intact rat brain metabolism measured by 13c nuclear magnetic resonance spectroscopy. J Neurosci 23, 5928– 5935 (2003).

56. J. G. Schulz et al., Glial β-oxidation regulates drosophila energy metabolism. Sci Rep 5, 7805 (2015).

57. M. Knobloch et al., A fatty acid oxidation-dependent metabolic shift regulates adult neural stem cell activity. Cell Rep 20, 2144–2155 (2017).

58. G. Jogl, Y. S. Hsiao, L. Tong, Structure and function of carnitine acyltransferases. Ann N Y Acad Sci 1033, 17–29 (2004).

59. P. A. Watkins, E. V. Ferrell, Jr., J. I. Pedersen, G. Hoefler, Peroxisomal fatty acid beta-oxidation in hepg2 cells. Arch Biochem Biophys 289, 329–336 (1991).

60. J. Demarquoy, F. Le Borgne, Crosstalk between mitochondria and peroxisomes. World J Biol Chem 6, 301–309 (2015).

61. M. Eisenberg-Bord, N. Shai, M. Schuldiner, M. Bohnert, A tether is a tether is a tether: Tethering at membrane contact sites. Developmental cell 39, 395–409 (2016).

62. G. Csordás, D. Weaver, G. Hajnóczky, Endoplasmic reticular-mitochondrial contactology: Structure and signaling functions. Trends Cell Biol, (2018).

63. O. M. de Brito, L. Scorrano, An intimate liaison: Spatial organization of the endoplasmic reticulum-mitochondria relationship. EMBO J 29, 2715–2723 (2010).

64. B. F. Lv et al., Protein tyrosine phosphatase interacting protein 51 (ptpip51) is a novel mitochondria protein with an n-terminal mitochondrial targeting sequence and induces apoptosis. Apoptosis 11, 1489–1501 (2006).

65. P. Koch et al., Expression profile of ptpip51 in mouse brain. J Comp Neurol 517, 892–905 (2009).

66. K. Kehrein et al., Organization of mitochondrial gene expression in two distinct ribosome-containing assemblies. Cell Rep, (2015).

67. M. Escobar-Henriques, T. Langer, Dynamic survey of mitochondria by ubiquitin. EMBO Rep 15, 231–243 (2014).

68. C. L. Chen et al., Proteomic mapping in live drosophila tissues using an engineered ascorbate peroxidase. Proc Natl Acad Sci U S A 112, 12093–12098 (2015).

69. D. Mahad, I. Ziabreva, H. Lassmann, D. Turnbull, Mitochondrial defects in acute multiple sclerosis lesions. Brain 131, 1722–1735 (2008).

70. I. J. Lodhi, C. F. Semenkovich, Peroxisomes: A nexus for lipid metabolism and cellular signaling. Cell Metab 19, 380–392 (2014).

71. A. Sugiura, S. Mattie, J. Prudent, H. M. McBride, Newly born peroxisomes are a hybrid of mitochondrial and er-derived pre-peroxisomes. Nature 542, 251–254 (2017).

72. P. Meera, S. M. Pulst, T. S. Otis, Cellular and circuit mechanisms underlying spinocerebellar ataxias. J Physiol 594, 4653–4660 (2016).

73. N. M. Márkus et al., Expression of mrna encoding mcu and other mitochondrial calcium regulatory genes depends on cell type, neuronal subtype, and ca2+ signaling. PloS one 11, e0148164 (2016).

74. J. Qiu et al., Mitochondrial calcium uniporter mcu controls excitotoxicity and is transcriptionally repressed by neuroprotective nuclear calcium signals. Nat Commun 4, 2034 (2013).

75. T. Zaglia et al., Content of mitochondrial calcium uniporter (mcu) in cardiomyocytes is regulated by microrna-1 in physiologic and pathologic hypertrophy. Proc Natl Acad Sci U S A 114, E9006–E9015 (2017).

76. Z. Dong et al., Mitochondrial ca2+ uniporter is a mitochondrial luminal redox sensor that augments mcu channel activity. Mol Cell 65, 1014–1028.e1017 (2017).

77. T. Henzi, B. Schwaller, Antagonistic regulation of parvalbumin expression and mitochondrial calcium handling capacity in renal epithelial cells. PloS one 10, e0142005 (2015).

78. M. Patron, H. G. Sprenger, T. Langer, M-aaa proteases, mitochondrial calcium homeostasis and neurodegeneration. Cell Res 28, 296–306 (2018).

79. V. Carelli, D. C. Chan, Mitochondrial DNA: Impacting central and peripheral nervous systems. Neuron 84, 1126–1142 (2014).

80. P. Mazzarelli et al., Carnitine palmitoyltransferase I in human carcinomas: A novel role in histone deacetylation? Cancer Biol Ther 6, 1606–1613 (2007).

